# Proteogenomic Analysis of Pancreatic Cancer Subtypes

**DOI:** 10.1101/2020.04.13.039834

**Authors:** Doris Kafita, Panji Nkhoma, Mildred Zulu, Musalula Sinkala

## Abstract

Pancreatic cancer remains a significant public health problem with an ever-rising incidence of disease. Cancers of the pancreas are characterised by various molecular aberrations, including changes in the proteomics and genomics landscape of the tumour cells. There is a need, therefore, to identify the proteomic landscape of pancreatic cancer and the specific genomic and molecular alterations associated with disease subtypes. Here, we carry out an integrative bioinformatics analysis of The Cancer Genome Atlas dataset that includes proteomics and whole-exome sequencing data collected from pancreatic cancer patients. We apply unsupervised clustering on the proteomics dataset to reveal the two distinct subtypes of pancreatic cancer. Using functional and pathway analysis, we demonstrate the different molecular processes and signalling aberrations of the pancreatic cancer subtypes. We explore the clinical characteristic of these subtypes to show differences in disease outcome. Using datasets of mutations and copy number alterations, we show that various signalling pathways are altered among pancreatic tumours, including the Wnt pathway, Notch pathway and PI3K-mTOR pathways. Altogether, we reveal the proteogenomic landscape of pancreatic cancer subtypes and the altered molecular processes which can be leveraged to devise more effective treatments.

## Introduction

Pancreatic cancer remains one of the deadliest malignancies [1,2]. Its incidence has continued to increase over the last few years predominantly because of life-style shifts in population and increase in life expectancy [3–6]. The molecular characterization of pancreatic cancer, including the transcriptomic, genomic, and epigenetic landscape has been studied by large-scale molecular profiling projects and many other studies [7–10]. Gene expression changes have led to the identification of molecular subtypes of the disease, which have the treatment and prognostic importance [11–13]. Also, many mutations and copy number changes are now known to drive pancreatic oncogenesis, and disease aggressiveness and alteration in various signal transduction pathways have now been identified [14–17].

Studies of many cancers including those of the breast, ovary and colon have shown that the transcriptome is poorly correlated to the proteome, with only 10%-20% of the variation in protein level explained by mRNA transcription levels [18,19]. Equally, no studies of pancreatic cancer have focused explicitly on segregating the patients based on the proteomic difference, and explored the subtype-specific alterations in the cellular processes and signalling pathways. Furthermore, recent studies have conducted a proteomic analysis of pancreatic cancer [20–26]; however, they have not integrated the proteomic subtypes with alterations in various signalling pathways, to the specific cancer gene mutations and/or the disease outcomes of the afflicted patients.

## Results

### Proteomics subtypes of pancreatic cancer

We applied unsupervised K-means clustering with a squared Euclidean metric to the proteomics data of the pancreatic cancer samples that are available in the TCGA to identify two consistent clusters of patient tumours (Figure 1a) [27]. These two clusters each comprised of 67 tumour samples that we named as subtype-1 and 34 tumour samples which we named as subtype-2. Furthermore, we evaluated the reproducibility of our two-cluster classification of pancreatic tumours using a supervised machine learning classification approach. Here, we used a Kernel naïve Bayes method [28] to show that we could accurately predict tumour subtypes with an accuracy of 98% (Figure 1b) and area under the curve of 0.97 (Figure 1c).

**Figure 1:**
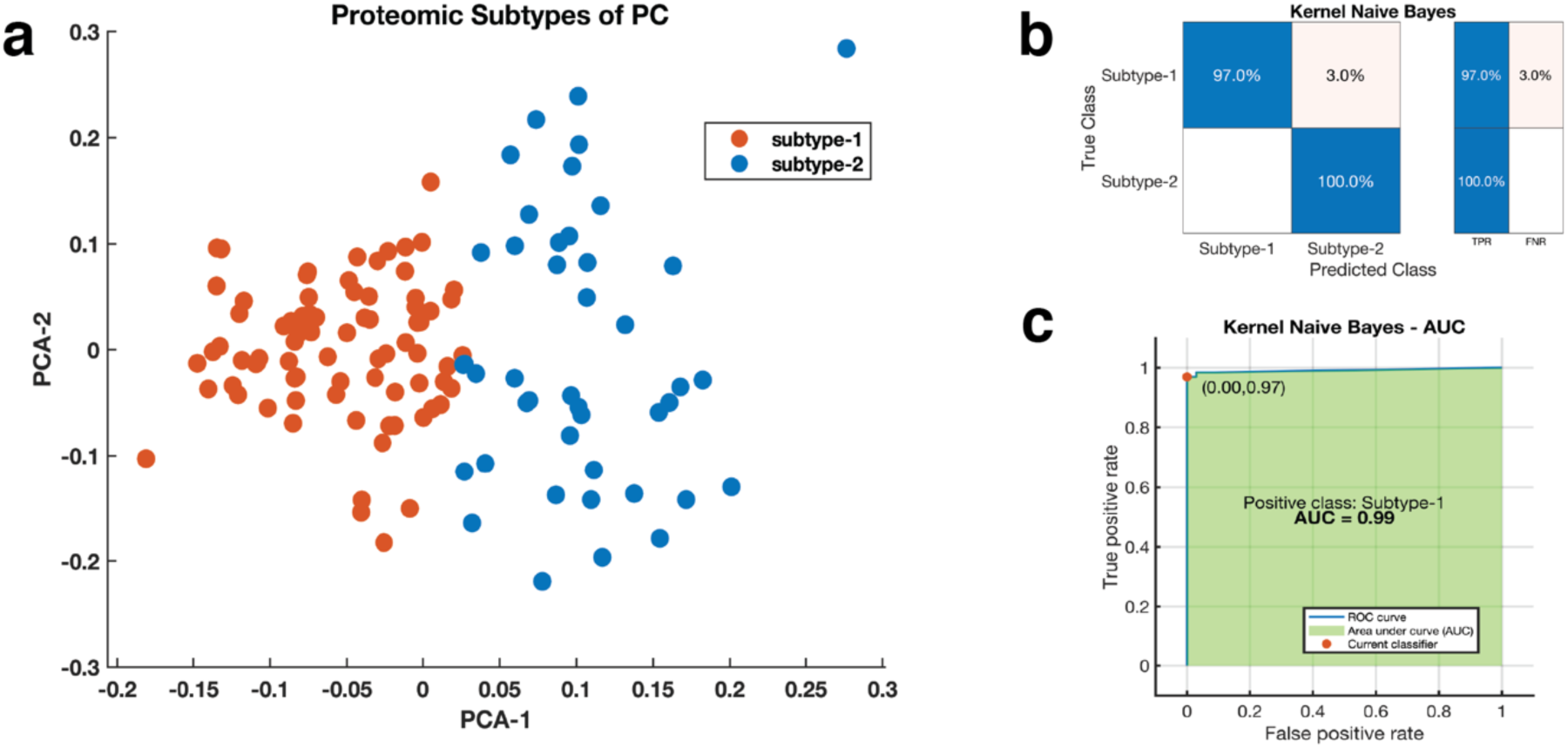
**(a)** Clustering of pancreatic tumours; the first and second principal components of a PCA analysis are plots on the x-axis and y-axis response. The points are coloured according to the K-mean clustering defined cluster assignments. **(b)** a representative confusion matrix for the Kernel naïve Bayes classifier used to validate the clustering of the proteomic subtypes of pancreatic cancer. The blue cells correspond to samples that are correctly classified. The red cells correspond to incorrectly classified samples. In the plot, TFR shows the true-positive rate and TNR indicate the false-negative rate. **(c)** the Receiver operating characteristic Curve for the Kernel naïve Bayes. The green shaded area represents the area under the curve (AUC = 0.99).

### Clinical characteristics of the proteomic subtypes of pancreatic cancer

We found a similar distribution of tumours of various grades (Figure 2a) and the patients who were living at the end of follow up (Figure 2b) for each of the two pancreatic cancer subtypes. Furthermore, we found similar distributions in the age, gender and the diagnosis of diabetes between the patients afflicted with the two subtypes of pancreatic cancer (Figure 2c). However, we showed that patients afflicted with subtype-2 tumours are significantly (χ^2^ = 10.8, p-value = 0.001) more progressive (57% of patients) compared to subtype-1 tumours (32%; Figure 2d).

**Figure 2:**
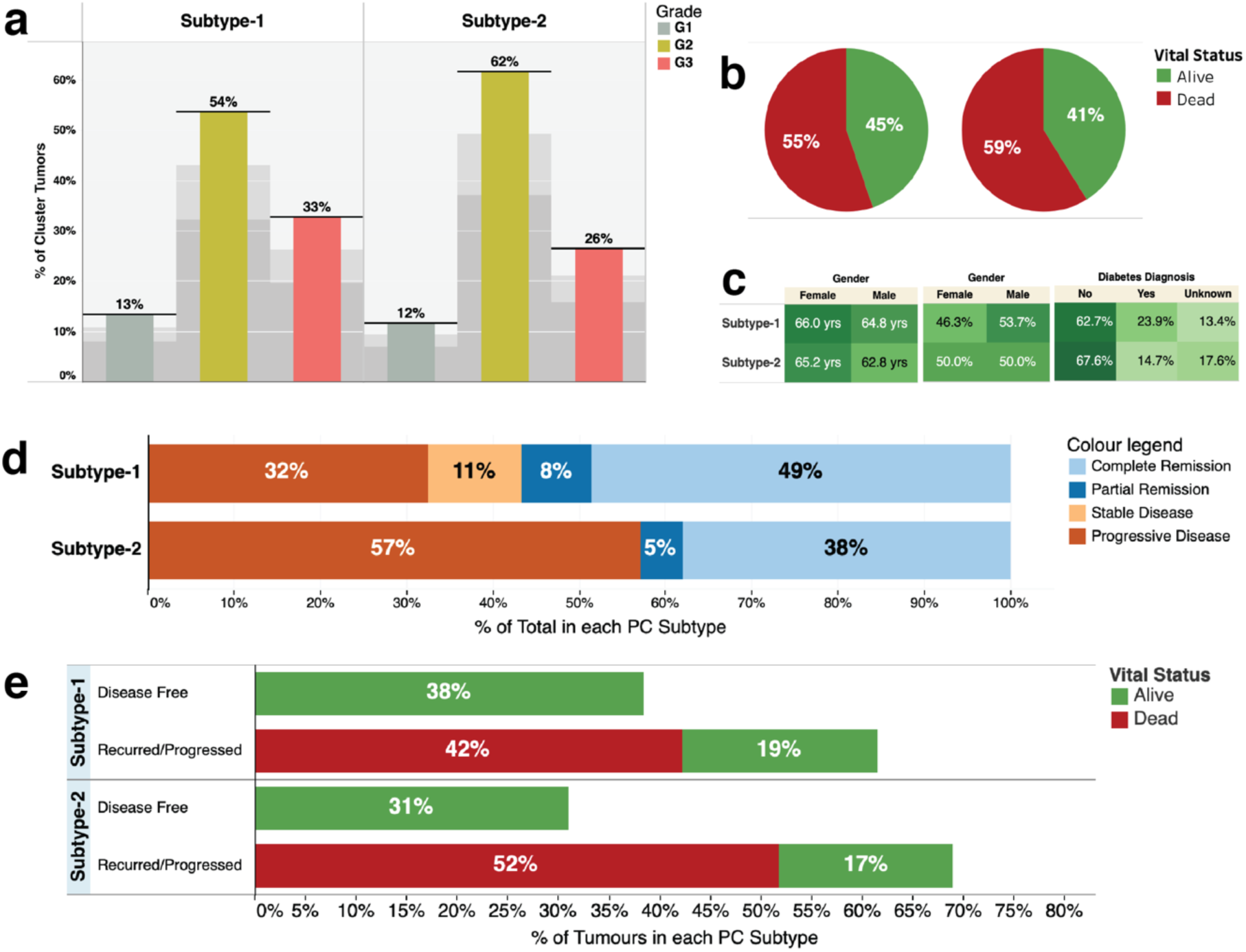
**(a)** Distribution of tumour grades across the proteomic subtypes: showing the percentage of the total count of the number of tumours for each grade of tumour broken down by proteomic subtype. **(b)** Pie chart showing the vital statistics after the first course of treatment across the two disease subtypes. **(c)** Highlight tables showing the distribution of, from left to right: the age of the study participants, the gender and the diabetes diagnosis. **(d)** The clinical outcomes after the first course of treatment across the disease subtypes. **(e)** Disease outcomes of each disease subtype. The bars are coloured based on vital statistics (dead or alive).

We found that of the patients with subtype-1 tumours, 38% were disease-free at the end of the follow-up, whereas 62% of the patients had a disease that either progressed or recurred, with only 19% of these patients surviving by the end of the follow-up period. For patients afflicted with subtype-2 tumours, 31% were disease-free at the end of the follow-up, whereas 57% of the patients had a disease that either progressed or recurred, with only 17% of these patients surviving by the end of the follow-up periods (Figure 2d). Overall, these findings show that subtype-2 tumours are more aggressive than subtype-1 tumours.

### Altered signalling pathways and molecular processes distinguish disease subtypes

Next, we compared the expression levels of various proteins between the two disease subtypes. We found that the subtype-1 tumours expressed significantly higher levels of several proteins, including mTOR, E-Cadherin, and Raf-pS338 when compared to the subtype-2 tumours (Supplementary File 1). Conversely, the subtype-2 tumours expressed significantly higher levels of several proteins including Stathmin, Mre11and MAP2K1 than the subtype-1 tumours (Supplementary File 1).

We compared the enrichment of KEGG pathways by querying Enrichr [29] with proteins that we found significantly up-regulated within tumours belonging to each disease subtype. Here, we found that some KEGG pathways that are altered in tumour belonging to both subtypes, albeit to different extents in some cases (Figure S1, also see Supplementary File 2).

We found that several KEGG pathways were significantly enriched for in only subtype-1 tumours. These included the mTOR signalling pathway (combined score [CS] = 1098, hypergeometric [HG] test p-value = 3.68 × 10^−14^), the ERBB2 signalling pathway (CS = 1625, HG p = 4.10 × 10^−14^) and among others (Figure S1a). Within the mTOR signalling pathway, we found several oncogenes, including mTOR, AKT1 and AKTS1 and tumour suppressor genes, including TSC1 and TSC2 and PTEN, that were significantly upregulated for subtype-1 tumours and all of which have previously been linked to pancreatic carcinogenesis (Supplementary File 1 and Supplementary File 2)[30–32].

Further, we found several KEGG pathways were significantly enriched for in subtypes-2 tumours but on in subtype-1 tumours. These included among others, the Hippo pathway (C.S. = 402, H.G. test p = 2.49 × 10^−7^) and the pathways associated with Chronic Myeloid Leukemia (C.S. = 817, H.G. test p = 4.3 × 10^−8^; Figure S2a and Supplementary File 2). These pathways encompass several known oncogenes (such as NRAS, CCND1, and ABL1) and tumour suppressor genes (such as SMAD4, CDKN1B and SMAD3) [33–35] (Figure 2C).

Next, we used the list of significantly up-regulated proteins within each disease subtypes to assessed for the molecular processes that drive the disease subtypes. Therefore, we queried Enrichr with the different protein lists described above to return the Gene Ontology (GO) Molecular Function terms that are enriched for in tumours of each disease subtype (see Supplementary File 2)[29]. For subtype-1 tumours, we found that the overexpressed proteins were enriched for, among others, the GO terms associated protein kinase activity (CS = 427, HG test p = 1.7 × 10^−12^) and protein serine/threonine/tyrosine kinase activity (CS = 1,707, HG test p = 1 × 10-5) (Figure 3a, also see Supplementary File 2). Conversely, for subtype-2 tumours, the proteins that we found up-regulated were enriched for, among others, GO terms that are associated with protein kinase binding (CS = 8.4, HG p = 2.2 × 10^−7^) and R-SMAD binding (CS = 305, HG p = 9.4 × 10^−4^; Figure 3b, also see Supplementary File 2).

**Figure 3:**
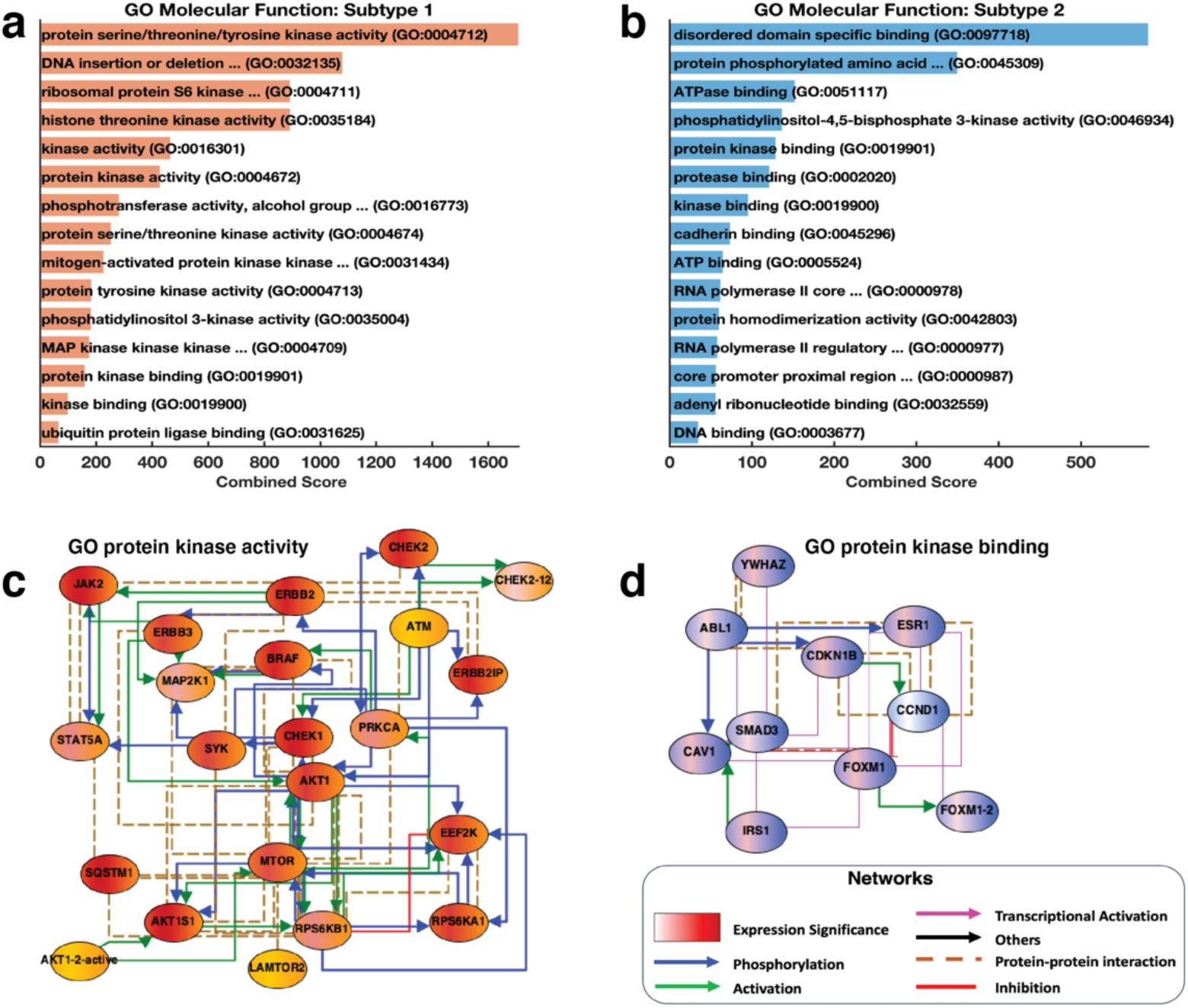
Gene ontology Molecular function: showing the top-ranked GO-terms molecular function enriched for **(a)** disease subtype-1 and **(b)** disease subtype-2 of pancreatic cancer based on the protein expression levels. **(c)** A network of the genes that encompass the GO-term molecular function “protein kinase activity” that we found highly enriched in subtype-1 tumours. The nodes are coloured using dual shades, orange for enrichment in the subtype-1 tumours and red represented the degree of statistical significance for each protein (negative logarithm of the p-values) between subtype-1 and subtype-2 tumours. **(d)** A network of the genes that encompass the GO-term molecular function “protein kinase binding” that we found highly enriched in subtype-2 tumours. The nodes are coloured using dual shades, blue for enrichment in the subtype-1 tumours and red represented the degree of statistical significance for each protein (negative logarithm of the p-values) between subtype-1 and subtype-2 tumours.

Overall, our findings revealed the distinct molecular mechanisms by which the development and progression of subtype-1 and subtype-2 may occur. For example, the GO term “protein kinase activity” forms a network whose nodes are significantly upregulated in subtypes-2 tumours, and are previously linked to oncogenesis, but in this case, we suggest that this likely only correct in subtype-2 tumours and not in subtype-1 tumours (Figure 3c). We also identified proteins involved in GO term “protein kinase binding”, a molecular process that our results suggest that would only play an essential role in the oncogenesis of the subtype-1 tumours (Figure 3d).

### The mutational landscape of proteomics subtypes of pancreatic cancer

We evaluated the extent of gene mutations and copy number variations (which we collectively refer to as gene alterations) in pancreatic cancer. Focusing only on the Consensus Cancer Genes [36], we found no significant difference in the gene alteration spectrum between the two pancreatic cancer subtypes (Figure 4a and Supplementary File 2). Across the disease subtypes, we revealed that, as previously reported by others, the most frequently altered genes were KRAS (altered in 91% of all samples), TP53 (71%), CDKN2A (42%) and SMAD4 (36%; Figure 4a) [2,12,37].

**Figure 4:**
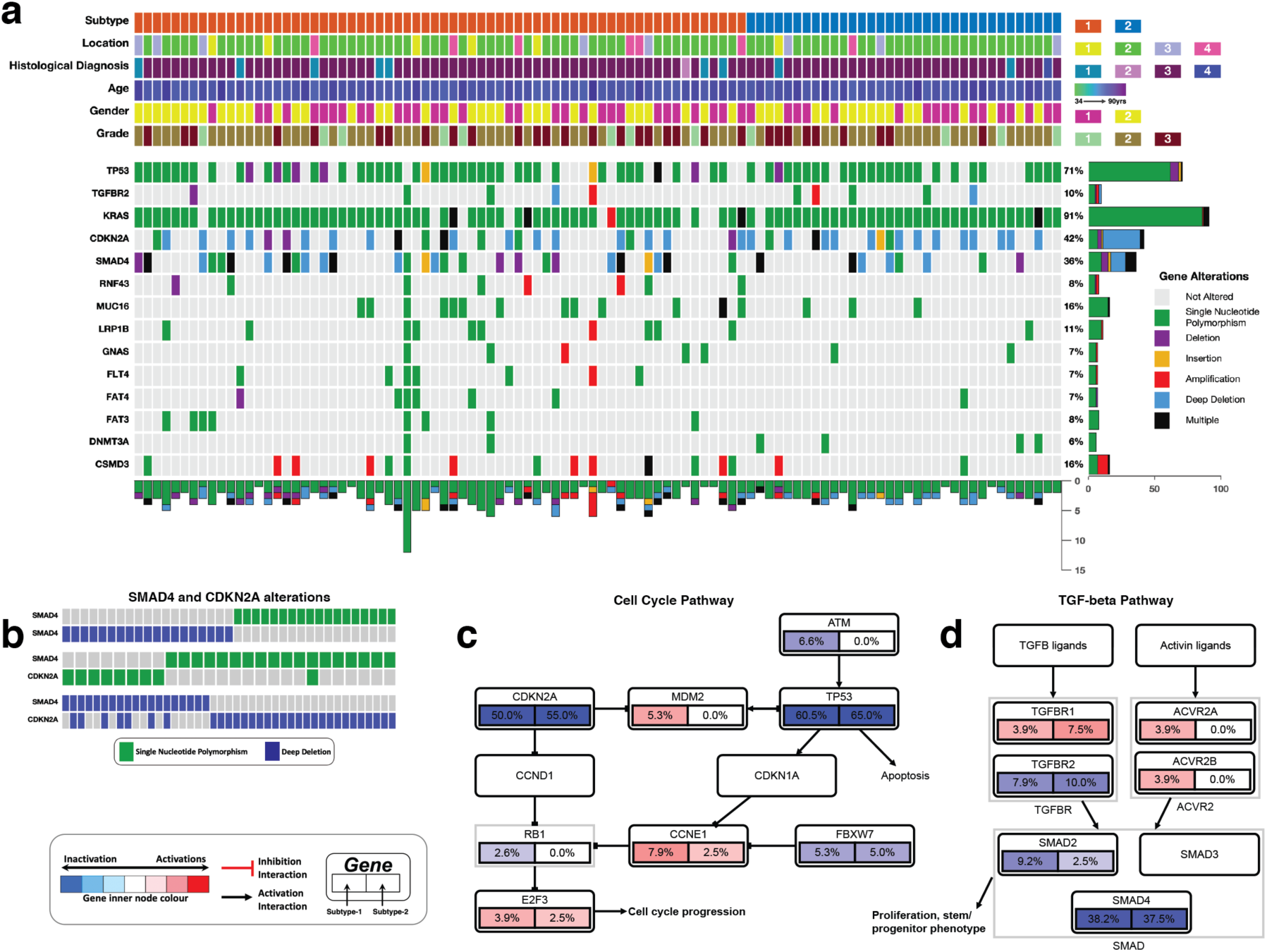
**(a)** The integrated plot of gene mutations, copy number alterations and the clinical features of the pancreatic tumours and the afflicted patients. From top to bottom panels indicate the proteomic subtypes of pancreatic cancer; the tumour location in the pancreas; the histological subtypes of the tumours; age at diagnosis; the patient’s gender; the tumour’s histological grade; non-silent mutations and copy number alteration frequency in each tumour across the altered genes. The key to the number coding of tumour location is 1; head, 2; body, 3; other, 4; tail. The number coding of histological diagnosis is 1; Pancreas-Adenocarcinoma-Other Subtype, 2; Pancreas-Colloid (mucinous non-cystic) Carcinoma, 3; Pancreatic Ductal Adenocarcinoma, 4; Discrepancy.

Interestingly, we observed that the mutations in SMAD4 were mutually exclusive to the copy number alterations in SMAD4 (Figure 4b). Likewise, we discovered that the mutations in SMAD4 and those in CDKN2A tended toward being mutual exclusivity (co-occurrence odd-radio = −0.137, p = 0.522). These findings show that the different gene alterations (either copy number alterations or mutations) in SMAD4 and CDKN2A drive pancreatic cells towards the malignant phenotype through perturbating the signalling pathways in which either SMAD4 or CDKN2A participate (Figure 4c and Figure 4d).

We found that in pancreatic tumours, genomic alterations are common within genes that are members of various well-known cell signalling pathways. Among the pathway with gene alterations were the Receptor tyrosine kinase – Ras pathway (altered in 75% of all tumours; Figure S2), Wnt pathway (altered in 29% of all tumours; Figure 5a), the PI3K-mTOR pathway (23%; Figure 5b) and Transforming Growth Factor Beta Signalling Pathway (48%; Figure 5c). These cell signalling pathways have been reported altered in various forms of cancers, including those of the lungs, skin, and breast, were they have also been shown to promote oncogenesis [15,38–40]. Accordingly, we suggest that these signalling pathways may play essential roles in pancreatic cancer and may present inflexion points for targeted therapies aimed at curing pancreatic cancer.

**Figure 5:**
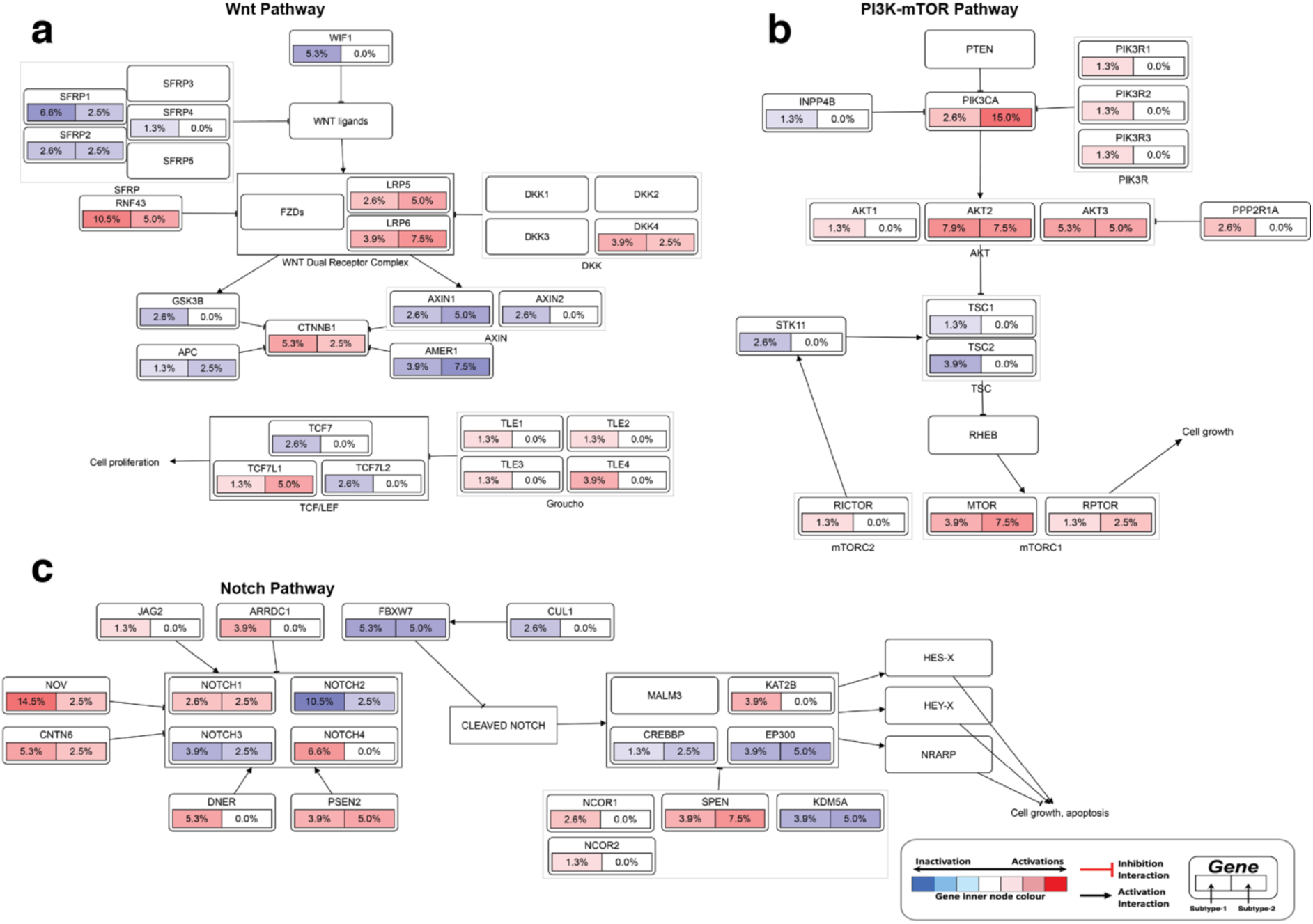
Alterations in **(a)** Wnt pathway, **(b)** PI3K-mTOR pathway and **(c)** Notch pathway. The node represents the percentage of each gene mutation and copy number alterations of (left half) subtype-1 and (right half) of subtype-2 pancreatic tumours. The nodes are coloured according to the types of genes: blue nodes for tumour suppressor genes and red for oncogenes. The interaction types are as given in the figure legend.

## Discussion

We conducted an integrated analysis of proteomics, clinical outcomes, mutations and copy number alterations of pancreatic cancer. Using machine learning, we showed that pancreatic tumours could be classified into two distinct subtypes. Patients afflicted of these disease subtypes show relatively similar demographics, suggesting that the onset of these cancer subtypes are not associated with the clinical parameters, such as age, diabetes, and gender. While the disease outcomes were vastly similar between the disease subtypes, we showed that patients afflicted with subtypes-2 were a significantly more progressive disease compared to those afflicted with subtypes-1 tumours.

We observed that subtype-1 tumours were significantly enriched for 28 distinct GO molecular functions, which is eight more than those that we found enriched for in the subtype-2 tumours (20 GO terms). Furthermore, the degree of dysregulation among the GO terms was on average higher for subtype-1 tumours (average C.S. = 416) than the subtype-2 tumours (average C.S. = 139). These findings are a divergence from those reported for other human cancers where it has been shown that the extent of signalling pathway perturbation is related to disease aggressiveness [41–43]. Here, we found that the subtype-2 tumours (the more progressive disease subtypes) are associated with fewer alterations to molecular processes.

Among the subtype-1 tumours, we found significant enrichment for kinase signalling events that are primarily downstream the ERBB2 and ERBB3 receptors (Figure 3c). Recent studies show that particular subtypes of pancreatic tumours respond well to anticancer agents that target the receptors tyrosine kinases of the epidermal growth factors signalling pathways [44,45]. Therefore, we expect that the subtypes-1 are possibly more responsive to drugs that target the ERBB class of receptors than will the subtype-2 tumours. Alternatively, we found that subtype-2 tumour showed enrichment for molecular functions in which various kinases participate, including the ABL1 kinase (Figure 3d). Accordingly, we suggest that, as has been shown with pancreatic tumours [46,47], subtype-2 are likely to be more responsive to drugs that target the ABL1 kinase and other related kinases.

Our results revealed that the mutations and copy number alteration were similar among the proteomic subtypes of pancreatic cancer. This finding may indicate that the primary genomic drivers of pancreatic oncogenes are similar among disease subtypes. The observed widespread alterations in the KRAS oncogene and TP53 tumour suppressor genes further confirm this (Figure 4a). As others have suggested, mutations in these genes likely perturb signalling through both the MAPK pathway, and p53 and cell cycle pathways [48,49]. Other gene alterations, such as those we found in the CDKN2A and SMAD4 genes (Figure 4b), which we have shown to be mutually exclusive, and among other gene alterations, should exert selective pressure that transforms the normal and pre-malignant cells [50–52]. Many other gene alterations in distinctive genes that participate in the same signalling pathway, such as the Notch pathway, PI3K-mTOR pathway (Figure S2) and metastasis pathway, likely come in later and contribute toward the progression of the disease [50,53].

Altogether, we have revealed the clinical and molecular characteristics of two distinct subtypes of pancreatic cancer. We have further shown that the altered signalling pathways and cellular processes differ between these disease subtypes, for which proteins of these pathways present a variety of potential disease subtype-specific biomarkers and drug targets.

## Methods

We analysed the TCGA project [43] datasets of 124 pancreatic cancer patients obtained from cBioPortal (http://www.cbioportal.org)[44]. Data on these patient samples included: reverse phase protein array-based (RPPA) proteomics data, D.N.A. copy number alterations and mutation data and comprehensive de-identified clinical and sample information.

### Proteomics classification of pancreatic cancer

We applied unsupervised machine learning methods to classify the pancreatic tumour samples based on the protein expression levels measured using RPPA. To evaluate the optimal number of clusters, we used the Calinski-Harabasz clustering evaluation criterion, which showed that the optimal number of clusters is two (Figure S3a) [45]. Then we applied unsupervised K-means over 1000 iterations with the squared Euclidean distance metric, and chose the clustering solution with highest average Silhouette score, to define the proteomic data to identify disease subtypes [46]. We visualised the clustering of the tumours; we reduced the dimensions of the proteomics measured data using Principal Component Analysis [47,48]. We plotted the first two dimensions of the principal components with points coloured based on the K-means clustering group assignment (Figure 1a). To evaluate the coherence of our K-mean clustering solution, we used an external validation method based on supervised re-classification of the tumours using a Kernel Naïve Bayes algorithm. Here, we used Bayesian optimisation to select the optimal machine learning hyperparameters for the Naïve Bayes algorithm (Figure S3b).

### Functional enrichment analyses

We identified the different proteins between the subtypes-1 and subtype-2 using the Welch t-test with the Benjamin-Hochberg adjustment for multiple comparisons applied to p-values [49,50]. Then, we used Enrichr to determine the KEGG pathway and GO molecular functions that are enriched for by the proteins that are up-regulated in each disease subtypes (see Supplement File 2) [29,51,52]. We compared the KEGG pathways and GO molecular functions between the two subtypes, to find those pathways and cellular process that are common and use to each disease subtypes.

Next, we retrieved known protein-protein interactions from the University of California Santa Cruz Super pathway, the Kinase Enrichment Analysis, and Chromatin Immunoprecipitation Enrichment Analysis [53–55]. Then, we used these interactions to connect proteins that are members of the most significantly enriched for GO molecular function terms for each disease subtypes. These GO-term were the protein kinase activity for subtype-1 tumours and protein kinase binding for the subtype-2 tumours. Then we used yEd to visualise the overall connectivity of two resulting networks (Figure 3c and Figure 3d).

### Analysis and mutations and copy number alterations

We evaluated the scope of gene alterations in the pancreatic subtypes using the mutation (single nucleotide polymorphisms and indels) data and copy number alterations data. We combined these two gene alteration data. Then we returned only genes that are associated with human cancers using information from the Sanger Consensus Cancer Gene Database [56]. Furthermore, the oncogenes and tumour suppressor genes in the gene alteration datasets were annotated using information from the UniProt Knowledgebase, the TSGene database, and the ONGene database [56–59]. We compared gene alterations between the disease subtypes using the Chi-square test. Also, we plotted the spectrum of genomic alterations for the fourteen most altered genes in the samples using a custom function (Figure 4a). To assess which signalling pathways, we used the PathwayMapper software [60].

### Statistical Analyses

We used MATLAB version 2020a to perform all the analyses presented here. The Fisher’s exact test was used to test for associations between categorical variables. The Welch test and the Wilcoxon rank-sum tests were used to compare differences in the tumours subtypes for the continuous variables among the various categories. We considered comparison as statistically significant when p-values are < 0.05 for single comparisons, and when the Benjamini-Hochberg adjusted p-values are < 0.05 for multiple comparisons.

## List of Abbreviations

TCGA: The Cancer Genome Atlas
RPPA: Reverse Phase Protein Array
GO: Gene Ontology
P.C.A.: Principle Component Analysis
H.G. test: Hypergeometric test
C.B.: combined score

## Declarations

### Author Contributions

The study was conceptualised by D.K.K., P.N., M.Z., and M.S.; The formal methodology was devised by D.K.K., P.N., M.Z., and M.S.; D.K.K., P.N. and M.S. performed the formal analysis of the datasets; D.K.K., M.Z., and P.N. wrote the draft manuscript; the manuscript was revised by was done D.K.K., P.N., M.Z., and M.S.; Visualisation we created M.S.

