## Supplementary Figures and Files for "Proteogenomic Analysis of Pancreatic Cancer Subtypes"

### Supplemental Information titles and legends

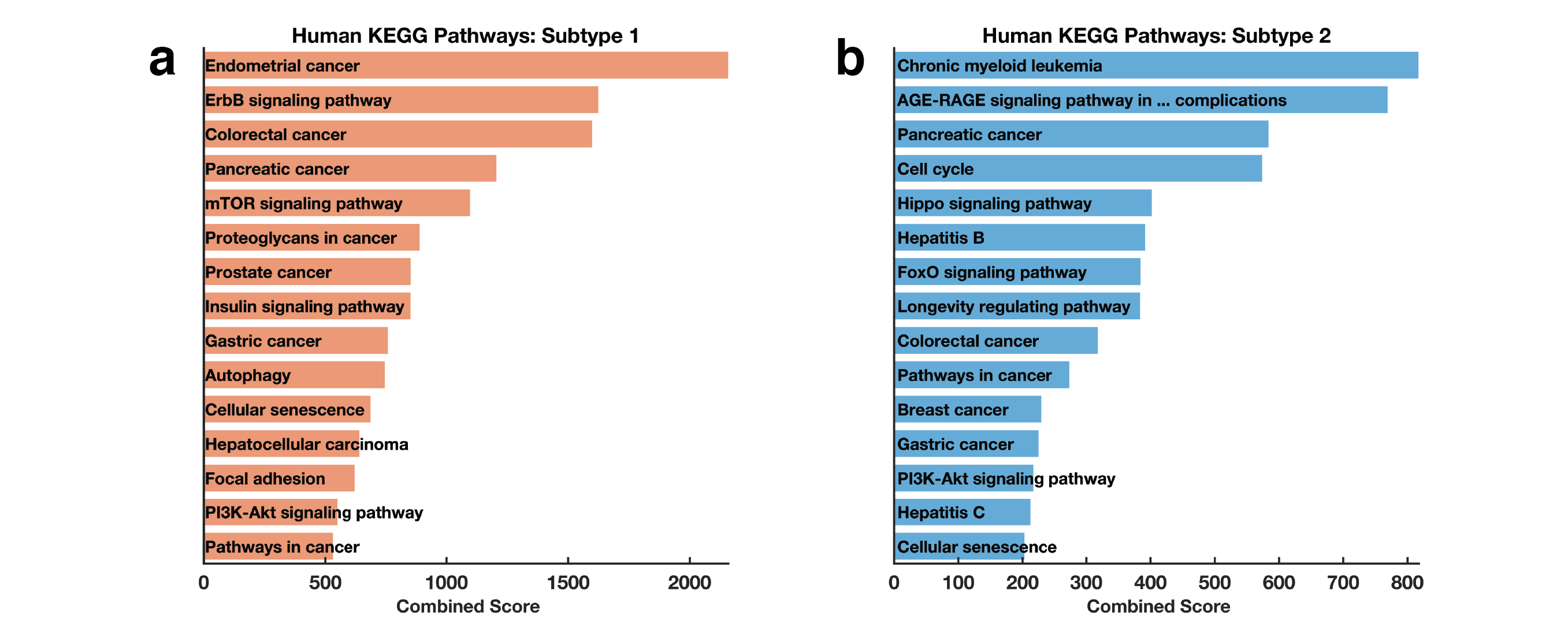

**Figure S1:** KEGG pathways: showing the top-ranked human KEGG pathway [1] enriched for **(a)** disease subtype-1 and **(b)** disease subtype-2 of pancreatic cancer based on the protein expression levels.

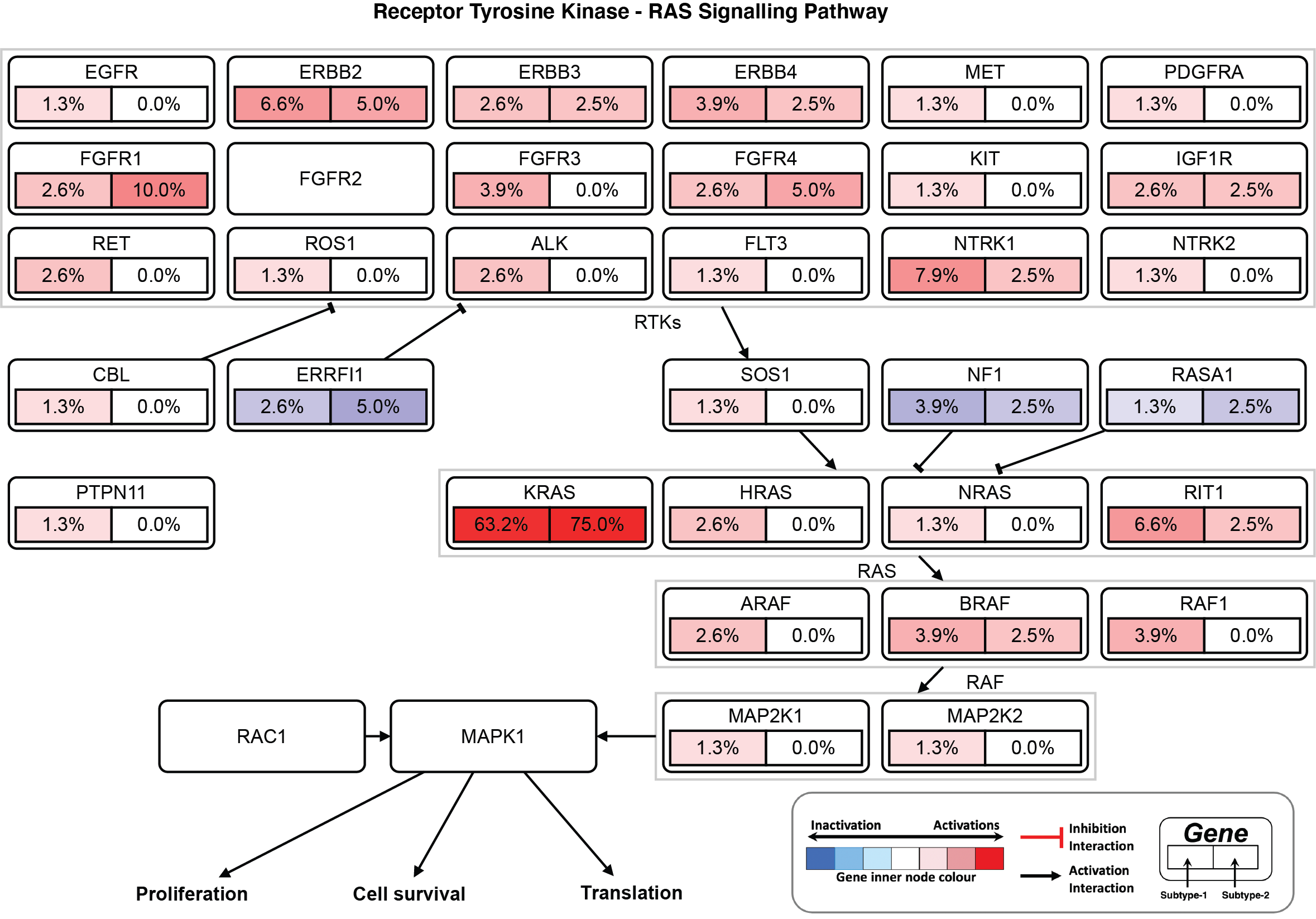

**Figure S2:** Alterations in Receptor Tyrosine Kinase – Rat Sarcoma (Ras) pathway. The node represents the percentage of each gene mutations and copy number alterations in (left half) subtype-1 and (right half) in subtype-2 pancreatic tumours. The nodes are coloured according to the types of genes: blue nodes for tumour suppressor genes and red for oncogenes. The interaction types are as given in the figure legend.

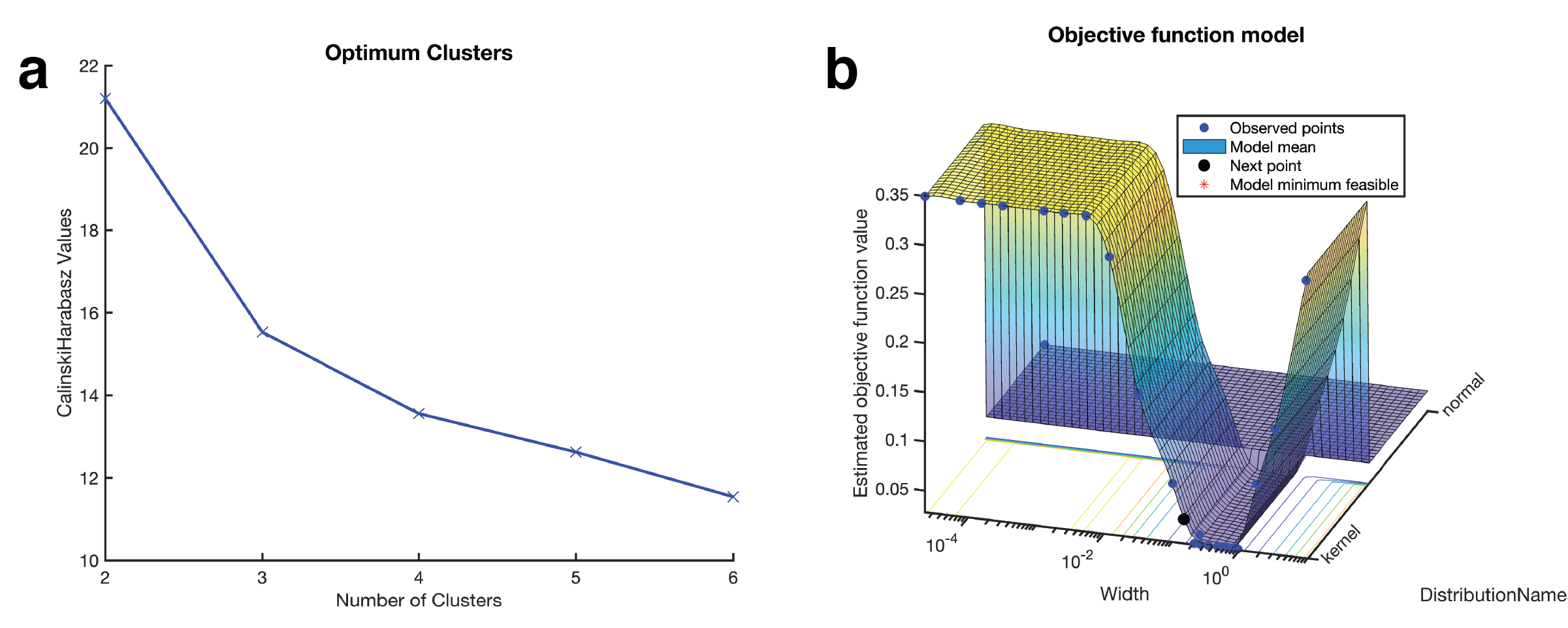

**Figure S3:** **(a)** Evaluating the optimum number of clusters: the plot displays the Calinski-Harabasz evaluation methods [2]. The optimum number of clusters is the number of cluster values that correspond to the highest Calinski-Harabasz value. In this case, the optimum number of clusters is two. **(b)** Range of values assessed by the Bayesian optimisation objective function to select the optimal machine learning hyperparameters for the Kernel naïve Bayes supervised learning model [3,4].

### Supplemental Files

**Supplementary File 1:** Differential expression results for proteins between subtype-1 and subtype-2 tumours of pancreatic cancer.

**Supplementary File 2:** KEGG pathways [1] and Gene Ontology [5] Molecular Function terms that are significantly enriched for in each subtype of pancreatic cancer.
